# *Candida auris* phenotypic heterogeneity determines pathogenicity *in vitro*

**DOI:** 10.1101/2020.04.20.052399

**Authors:** Jason L Brown, Chris Delaney, Bryn Short, Mark C Butcher, Emily McKloud, Craig Williams, Ryan Kean, Gordon Ramage

**Affiliations:** Glasgow Biofilm Research Network, Glasgow Dental School, School of Medicine, College of Medical, Veterinary and Life Sciences, University of Glasgow, Glasgow, UK, G12 8TA; Department of Life Sciences, School of Health and Life Sciences, Glasgow Caledonian University, Glasgow, UK, G4 0BA; Microbiology Department, Lancaster Royal Infirmary & University of Lancaster

**Keywords:** *Candida auris*, aggregate, host-pathogen interactions, *in vitro* skin model

## Abstract

*Candida auris* is an enigmatic yeast that provides substantial global risk in healthcare facilities and intensive care units. A unique phenotype exhibited by certain isolates of *C. auris* is their ability to form small clusters of cells known as aggregates, which have been to a limited extent described in the context of pathogenic traits. In this study, we screened several non-aggregative and aggregative *C. auris* isolates for biofilm formation, where we observed a level of heterogeneity amongst the different phenotypes. Next, we utilised an RNA-sequencing approach to investigate the transcriptional responses during biofilm formation of a non-aggregative and aggregative isolate of the initial pool. Observations from these analyses indicate unique transcriptional profiles in the two isolates, with several genes identified relating to proteins involved in adhesion and invasion of the host in other fungal species. From these findings we investigated for the first time the fungal recognition and inflammatory responses of a three-dimensional skin epithelial model to these isolates. In these models, a wound was induced to mimic a portal of entry for *C. auris*. We show both phenotypes elicited minimal response in the model minus induction of the wound, yet in the wounded tissue both phenotypes induced a greater response, with the aggregative isolate more pro-inflammatory. This capacity of aggregative *C. auris* biofilms to generate such responses in the wounded skin highlights how this opportunistic yeast is a high risk within the intensive care environment where susceptible patients have multiple indwelling lines.

**Importance:** *Candida auris* has recently emerged as an important cause of concern within healthcare environments due to its ability to persist and tolerate commonly used antiseptics and disinfectants, particularly when surface attached (biofilms). This yeast is able to colonise and subsequently infect patients, particularly those that are critically ill or immunosuppressed, which may result in death. We have undertaken analysis on two different types of this yeast, using molecular and immunological tools to determine whether either of these has a greater ability to cause serious infections. We describe that both isolates exhibit largely different transcriptional profiles during biofilm development. Finally, we show that the inability to form small aggregates (or clusters) of cells has an adverse effect on the organisms immuno-stimulatory properties, suggestive the non-aggregative phenotype may exhibit a certain level of immune evasion.

## Introduction

*C. auris* is a nosocomial pathogen first identified in 2009 [1]. To date, this multidrug resistant organism has been identified in over 40 countries on 6 different continents, providing a substantial global risk in healthcare facilities and intensive care units [2–4]. It is postulated that the emergence of *C. auris* may have coincided with climate change based on its particular attributes, resulting in an thermotolerant organism with the ability to persist in the environment before transmission to humans [5].

A unique pathogenic trait exhibited by some isolates of *C. auris* is their ability to form aggregates (Agg) [6–8]. Despite the well-documented prevalence of *C. auris* worldwide, relatively little is known about the Agg phenotype of the organism. The existence of four geographically and phylogenetically distinct clades of the organism [2], and a fifth recently proposed [9], has restricted a definitive profiling of *C. auris* pathogenic mechanism of these aggregates in regard to biofilm forming capabilities, drug resistance pathways and interactions with the host. Of the publications that exist, these have documented characteristic pathogenic traits for both phenotypes *in vitro* and *in vivo* [6, 10, 11]. Others have shown that the Agg phenotype is inducible under certain conditions [7, 8], whilst histological analyses of murine models have shown that aggregates can accumulate in organs following *C. auris* infection [7, 12, 13]. Therefore, further studies are required to investigate this characteristic Agg phenomenon in *C. auris* isolates to fully comprehend the pathogenic pathways of the organism, and to understand how such mechanisms may differ from their non-Agg counterparts.

Limited evidence also exists for studies investigating the interactions of *C. auris* with components of the host, although several *in vivo* models have been employed to document the virulence of *C. auris*. Of these, *Galleria mellonella* larvae infection models and murine models of invasive candidiasis have shown varying survival rates post-infection with *C. auris* [6, 8, 10, 12–15], reaffirming that genetic variability amongst clades impacts the organisms virulence. However, such studies have been limited in investigating the host immune response to the organism. Recently, Johnson et al (2019) utilised a Zebrafish (*Danio rerio*) model to monitor *C. auris*-host cell interactions *in vivo*. This work highlighted that *C. auris* (strain B11203 Indian isolate phylogenetically part of the South Asian or India/Pakistan clade [2], which appeared to exhibit a non-Agg phenotype) was resistant to neutrophil-mediated killing, suggesting that the organism has the ability to persist incognito in the host [16]. *In vitro* studies have shown interactions between *C. auris* and epithelial tissue, emphasizing that the organism can persist on skin. In this study, Horton and colleagues (2020) demonstrated that *C. auris* (B11203 strain, as above) formed high-burden biofilms on porcine skin biopsies in the presence of an artificial sweat medium [17]. To date, no studies have investigated the host inflammatory response to the non-Agg and/or Agg phenotype.

In this study, we sought to investigate the level of heterogeneity amongst different non-Agg and Agg isolates. We deemed this pertinent given that such traits of heterogeneity amongst isolates have previously been described for other *Candida* species, significantly impacting clinical outcomes and mortality rates [18]. To further investigate this Agg versus non-Agg phenotype, transcriptome analyses were performed on planktonic cells and biofilms of two selected isolates from the initial pool. Upon completion of these analyses, we discovered that several genes associated with cell membrane and/or cell wall proteins (e.g. cellular components) were upregulated in the Agg biofilm. Such unique transcriptional profiles in respect to the cellular components led us to investigate the host response following stimulation with both *C. auris* phenotypes *in vitro*. For this, a two- and three-dimensional skin wound model was employed to investigate the epithelial response to the Agg and non-Agg isolates of *C. auris*. Both skin wound models exhibited different profiles to both isolates indicating unique fungal recognition and/or host response to the Agg and non-Agg phenotype. Interestingly, there was minimal response by the host to *C. auris* without induction of the wound, suggestive the organism relies on loss of tissue integrity to become invasive.

## Methods

### Microbial growth and standardisation

For *in vitro* biofilm biomass assessment, a pool of aggregating (Agg; n=12) and single-celled, non-aggregative (non-Agg; n=14) *C. auris* clinical isolates (gifted by Dr Andrew Borman and Dr Elizabeth Johnson, Public Health England, UK). All *C. auris* isolates were stored in Microbank™ beads (Pro-lab Diagnostics, UK) prior to use. Each isolate was grown on Sabouraud dextrose (SAB) agar (Oxoid, UK) at 30°C for 24-48 h then stored at 4°C prior to propagation in yeast peptone dextrose (YPD; Sigma-Aldrich, UK) medium overnight (16 h) at 30°C, gently shaking at 200 rpm. Cells were pelleted by centrifugation (3,000 x g) then washed two times in phosphate buffered saline (PBS). Cells were then standardised to desired concentration following counting using a haemocytometer, then resuspended in selected media for each assay, as described within.

*C. auris* isolate phenotypes were determined visually by suspending one colony in 1 mL (PBS, Sigma-Aldrich, UK). Isolates were termed as ‘aggregators’ if the added colony did not disperse upon mixing in PBS. For RNA sequencing and transcriptional analysis of *C. auris* biofilms and co-culture systems, one Agg (NCPF 8978) and non-Agg (NCPF 8973) isolate was used.

### Biofilm growth and biomass assessment

Fungal cells were adjusted to 1 × 10^6^ cells/mL in Roswell Parks Memorial Institute (RPMI) media (Sigma-Aldrich, UK) and biofilms formed for 4 or 24 hours at 37°C in flat-bottom wells of 96-well plates (Corning, UK). Appropriate media controls were included on each plate to test for contamination. Following incubation, biofilms were washed gently once in PBS to remove any non-adhered cells. The biomass of each biofilm was determined via 0.05% crystal violet (CV) staining as described previously [19]. Absorbance of the CV stain was measured spectrophotometrically at 570nm in a microtiter plate reader (FLUOStar Omega, BMG Labtech, UK).

### Monitoring the growth of Candida auris biofilms in real time

The xCELLigence real time cell analyser (RTCA, ACEA Bioscience Inc, San Diego, CA) was used to monitor the formation of *C. auris* biofilms in real time using electron impedance measurements (presented as cell index, CI) which is directly related to cell attachment and proliferation. In brief, the E-plate containing 100 μL of pre-heated RPMI medium was loaded into the RTCA which had been placed in the incubator 2 h prior to the experiment to test media impedance and electrode connectivity. Cultures of each *C. auris* isolate used in this study were standardised to 2 × 10^6^ CFU/mL and added to the E-plate in 100 μL aliquots in triplicate. Appropriate media controls minus inoculum were also included in triplicate. Biofilm formation was measured over 24 hours with CI readings taken every 5 minutes. Normalised CI values were exported from the RTCA software and analysed in GraphPad Prism (version 8; GraphPad Software Inc., La Jolla, CA). More detailed descriptions for this technology can be found elsewhere [20].

### Two-dimensional monolayer co-culture model

Adult human epidermal keratinocytes (HEKa) cells (Invitrogen, GIBCO®, UK) were used for two-dimensional co-culture experiments. Frozen stocks of HEKa cells (1 × 10^6^ cells/ml; passage number lower than 10) were revived and seeded in T-75 tissue culture flasks (Corning, UK) in media 154 (Thermo-Fisher, UK) supplemented with 100 U/ml of pen/strep and human keratinocyte growth supplement (HKGS) (Thermo-Fisher, UK). The flasks were incubated at 37°C (5 % CO_2_) and media was changed every 48 h until the cells reached 80-90% confluence [21]. Confluent cells were passaged using 0.05% trypsin EDTA (Sigma-Aldrich, UK) and the enzymatic reaction inhibited using trypsin neutralizer solution (Sigma-Aldrich, UK). Passaged cells were then seeded into 24-well plates (Corning, UK) at final concentration 2 × 10^5^ cells/ml. After 24 to 48 h the cells reached the adequate confluence for *C. auris* co-culture experiments as described below.

### Three-dimensional human epidermis co-culture model

Reconstituted human epidermis (RHE) used for 3D co-culture experiments was purchased from Episkin (Skin Ethic™, Lyon, France http://www.episkin.com). RHE was formed from normal human keratinocytes cultured on an inert polycarbonate filter at the air-liquid interface, in a chemically defined medium grown to 17-day maturity. This model is histologically similar to *in vivo* human epidermis. Upon arrival and prior to experimental set-up, RHE was incubated with maintenance media in 24-well plates (Corning, UK) for 24 h, 5% CO_2_ at 37°C. Maintenance media was replaced then the co-culture three-dimensional system was set up as described below.

### Wound model in two-dimensional and three-dimensional co-culture systems

HEKa cell monolayers were scratched using a similar method to as previously described to mimic a wound model [22, 23]. Briefly, monolayers were grown to confluence as described above then three parallel scratches were introduced across the surface using a 100 μL pipette tip prior to inoculation with *C. auris*. For RHE, a 19-gauge needle was used to scratch the tissue. For all co-culture experiments, Agg *C. auris* NCPF 8978 and non-Agg *C. auris* NCPF 8973 were grown as described above, then standardised to 2 × 10^6^/mL (multiplicity of infection of 10 to HEKa cells; MOI 10, and as previously described for *Candida*-tissue co-culture [24]). For the two-dimensional system, 2 × 10^6^/mL *C. auris* was prepared in 500μL supplemented media 154 and added directly to the confluent HEKa cells. For the three-dimensional co-culture model, 2 × 10^6^/mL *C. auris* was prepared in 100μL of sterile PBS and this suspension was added directly to the RHE tissue. All experiments were conducted for 24 hr at 5% CO_2_. Infected non-scratched HEKa cells and RHE tissue were used as controls e.g., no wound, and uninoculated co-cultured cells or tissues were also included for all experiments. All control and wounded models were infected in triplicate with both isolates of *C. auris*.

### Histological staining of skin epidermis

Following co-culture with *C. auris*, epithelial tissue was carefully cut from the 0.5 cm^2^ insert using a 19-gauge needle and washed three times in sterile PBS to remove non-adherent cells in a similar manner as previously described [25], as summarized in the schematic in Figure 1. Tissue was then fixed in 10% neutral-buffered formalin prior to embedding in paraffin. A Finnesse ME+ microtome (Thermo Scientific, UK) was used to cut 2 μm sections and tissue sections stained with haematoxylin and eosin, or with the fungal specific Periodic acid-Schiff (PAS) reagents and counter-stained with haematoxylin.

**Figure 1.**
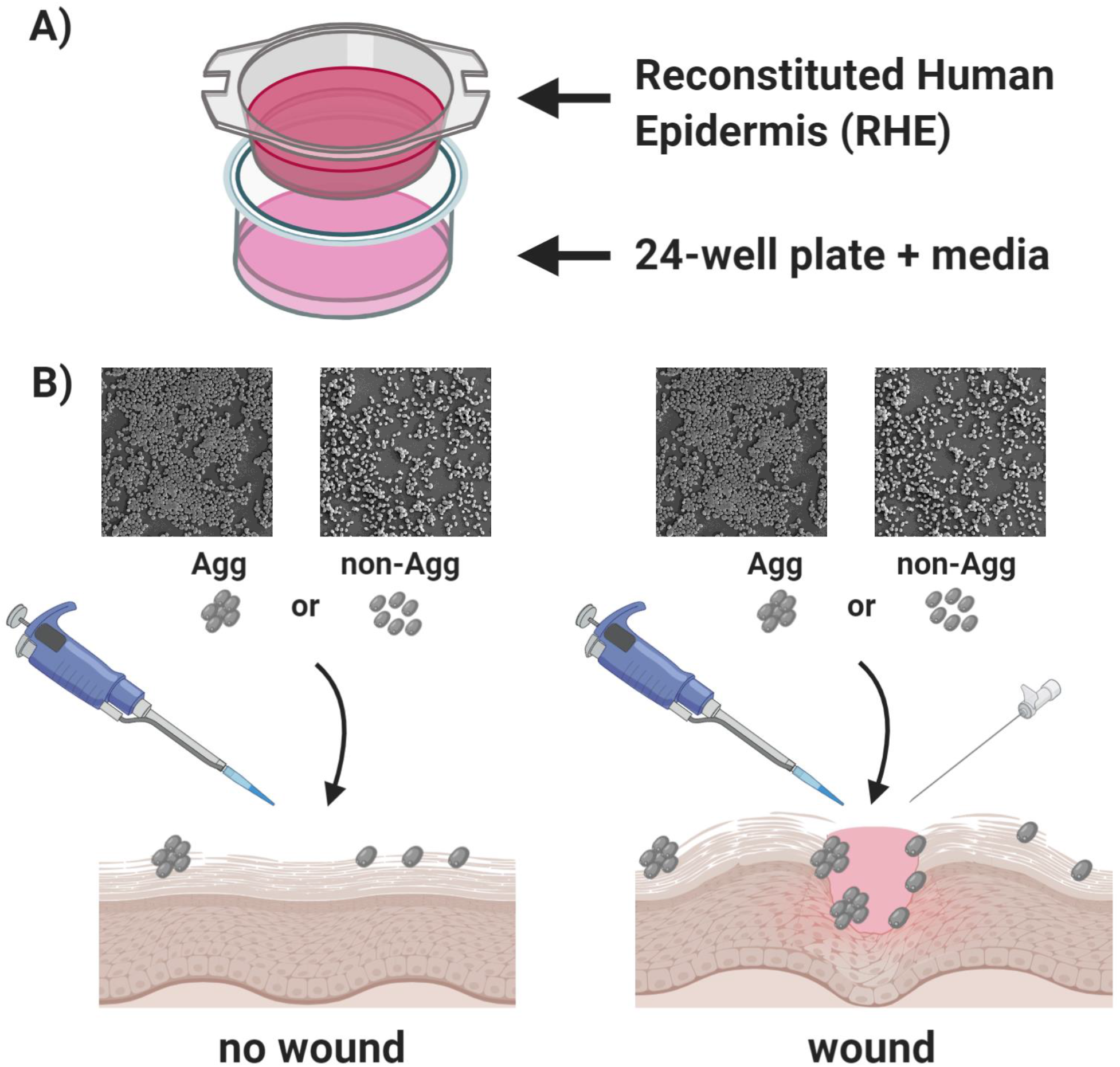
Schematic diagram depicting the experimental set up for the three-dimensional co culture of skin epidermis and *Candida auris*. A 17-day mature reconstructed human epidermis (RHE) on 0.5 cm^2^ inserts was purchased from Episkin (Skin Ethic™). Inserts were carefully lowered into 24-well plates containing maintenance media supplied by the company (panel A). To assess the host response to aggregative and non-aggregative *C. auris*, control and wounded tissue was co-cultured with both isolates (NCPF 8973 and NCPF 8978) (panel B). A total of 2 × 10^6^ fungal cells in 100 μL PBS was added to the tissue and incubated overnight at 37°C, 5% CO_2_. For some tissues, prior to addition of fungal inoculum, three scratch wounds were induced using a sterile 19 G needle across the surface of the tissue. For visual representation of phenotype, scanning electron microscopy images included in panel B from 24 h biofilms, clearly showing the differences in cellular phenotypes between the two *C. auris* isolates. These images were taken at x 1000 magnification as viewed under a JEOL JSM-6400 scanning electron microscope (samples processed as previously described [66]).

### Epithelial cell viability

To assess any cytotoxic effects of *C. auris* on HEKa cells and RHE tissue, a Pierce lactate dehydrogenase (LDH) Cytotoxicity Assay Kit (Thermo Scientific; UK) was used according to the manufacturers’ instructions. Following co-culture, cell or tissue spent media was assayed using the above kit to quantify the level of LDH release as a measure of host cellular disruption.

### Differential gene expression analysis

HEKa cells and RHE tissue following co-culture were lysed in RLT lysis buffer (Qiagen Ltd, UK) containing 0.01% (v/v) β-2-mercaptoethanol (β2ME) before bead beating. All RNA was extracted using the RNeasy Mini Kit according to manufacturers’ instructions (Qiagen Ltd, UK) and quantified using a NanoDrop 1000 spectrophotometer [Thermo Scientific, UK]. RNA was converted to complementary DNA (cDNA) using the High Capacity RNA to cDNA kit (Life Technologies, UK) as per the manufacturer’s instructions. Gene expression was assessed using SYBR Green^ER^ based-quantitative polymerase chain reaction (qPCR) or RT^2^ profiler arrays (Qiagen Ltd, UK). For SYBR Green^ER^ based-qPCR analyses, the following PCR thermal profiles was used; holding stage at 50°C for 2 minutes, followed by denaturation stage at 95°C for 10 minutes and then 40 cycles of 95°C for 3 seconds and 60°C for 15 seconds. qPCR plates were run on the StepOnePlus™ Real-Time PCR System. The following primer sequences were used for SYBR Green^ER^ based-qPCR analyses of host cells; *GAPDH*, forward primer, 5’ to 3’, CAAGGCTGAGAACGGGAAG and reverse primer, 5’ to 3’, GGTGGTGAAGACGCCAGT [26]; *IL8*, forward primer, 5’ to 3’, CAGAGACAGCAGAGCACACAA and reverse primer, 5’ to 3’, TTAGCACTCCTTGGCAAAAC [27]. For gene expression analyses of *C. auris*, primers for adhesin gene *ALS5* and proteinase gene *SAP5* were used as follows; *ALS5;* forward primer, 5’ to 3’, ATACCAGGGTCGGTAGCAGT and reverse primer, 5’ to 3’, CTATCTTCGCCGCTTGGGAT and *SAP5*; forward primer, 5’ to 3’, GGATGCAGCTCTTCCTGGTT and reverse primer, 5’ to 3’, CTTCCAGTTTGCGGTTGTGG. For other gene expression analyses of RHE tissue, a custom designed RT^2^ Profiler PCR array was compiled containing primers for genes associated with inflammation and fungal recognition or stimulation of host tissue. For these arrays, the following thermal cycle was used on the MxProP Quantitative PCR machine; 10 minutes at 95°C followed by 40 cycles of 15 seconds at 95°C and 60 seconds at 60°C, and data assembled using MxProP 3000 software (Stratagene, Netherlands). Expression levels for all genes of interest were normalized to the housekeeping gene, *β-actin* for *C. auris* gene expression and *GAPDH* for mammalian cells, according to the 2-ΔCt method, and then quantified using the 2-ΔΔCt method [28].

### RNA sequencing

For RNA sequencing of *C. auris* biofilms, RNA was extracted from 24-h *C. auris* biofilms as described previously [29]. In brief, biofilms were grown as above on Thermanox coverslips (Thermo-Fisher, UK) in 24 well plates (Corning, UK). Biofilms were removed from coverslips by sonication at 35 kHz for 10 minutes in a sonic bath in 1 ml of PBS and the sonicate transferred to a 2.0 ml RNase-free bead beating tube (Sigma-Aldrich, UK). Cells were homogenized in TRIzol ™ (Invitrogen, UK) with 0.5 mm glass beads using a BeadBug microtube homogeniser for a total of 90 seconds (Benchmark-Scientific, USA). RNA was then extracted as described above using the RNeasy Mini Kit according to manufacturers’ instructions (Qiagen Ltd, UK). Following extraction, RNA quality and quantity were determined using a Bioanalyzer (Agilent, USA), where a minimum RNA integrity number and quantity of 7 and 2.5 μg, respectively were obtained for each sample. Annotation of data following sample submission to Edinburgh Genomics (http://genomics.ed.ac.uk/) was completed as previously described [30]. Briefly, raw fastq reads were trimmed and aligned to the *Candida auris* representative genome B8441 using Hisat2 [31]. Reads were then processed and assembled *de novo* using the Trinity assembly pipeline [32]. Trinotate and Blast2Go were utilised to assign gene IDs to homologous sequences using BLAST and Interpro [33, 34]. Differential expression analysis was performed according to DESeq2 pipeline and functional over representation was determined using GOseq [35, 36] within R. Additionally visualisation of overrepresented pathways were drawn within R. Raw data files for these analyses are deposited under accession no. PRJNA477447 (https://www.ncbi.nlm.nih.gov/bioproject/477447).

### Statistical analysis

Statistical analyses were performed using GraphPad Prism. Two-tailed paired or unpaired Student’s t-tests were used to compare the means of two samples as stated within, or one-way analysis of variance (ANOVA) to compare the means of more than two samples. Tukey’s post-test was applied to the p value to account for multiple comparisons of the data. P-values of <0.05 were considered statistically significant.

## Results

The Agg phenotype is a unique trait of *C. auris,* one that can influences the organisms pathogenic traits *in vitro* and *in vivo* [6, 10, 11]. To corroborate these previous observations, differences in early and late biofilm formation were assessed between non-Agg and Agg isolates of *C. auris*. A total of 26 non-Agg and Agg *C. auris* clinical isolates were screened during early (4 h) and mature (24 h) biofilm growth stages (Figure 2A and B). Both sets of non-Agg and Agg *C. auris* isolates formed biofilms in a time-dependant manner as assessed by crystal violet staining. With the exception of NCPF 8993 (non-Agg) and NCPF 8996 (Agg), all isolates formed biofilms with greater biomass after 24 hours culture compared to 4 hours (Figure 2C and D). Although, the Agg phenotype tended to form biofilms with greater biomass at both 4 h and 24 h, we observed no statistical differences when collectively comparing all non-aggregating and aggregating isolates of *C. auris* at either timepoint (Supplementary Figure 1A and B). From this data, it was clear that some isolates formed biofilms with greater biomass than others, suggestive of a certain level of heterogeneity between isolates of Agg and non-Agg phenotype, in line with previous observations [6, 10, 37]. Similar trends of biofilm heterogeneity were observed when monitoring biofilm formation via impedance measurements in real time at both timepoints in non-Agg and Agg phenotype (Figure 2E-H and Supplementary 1C and D). These data support corroboration of these unique phenotypes.

**Figure 2.**
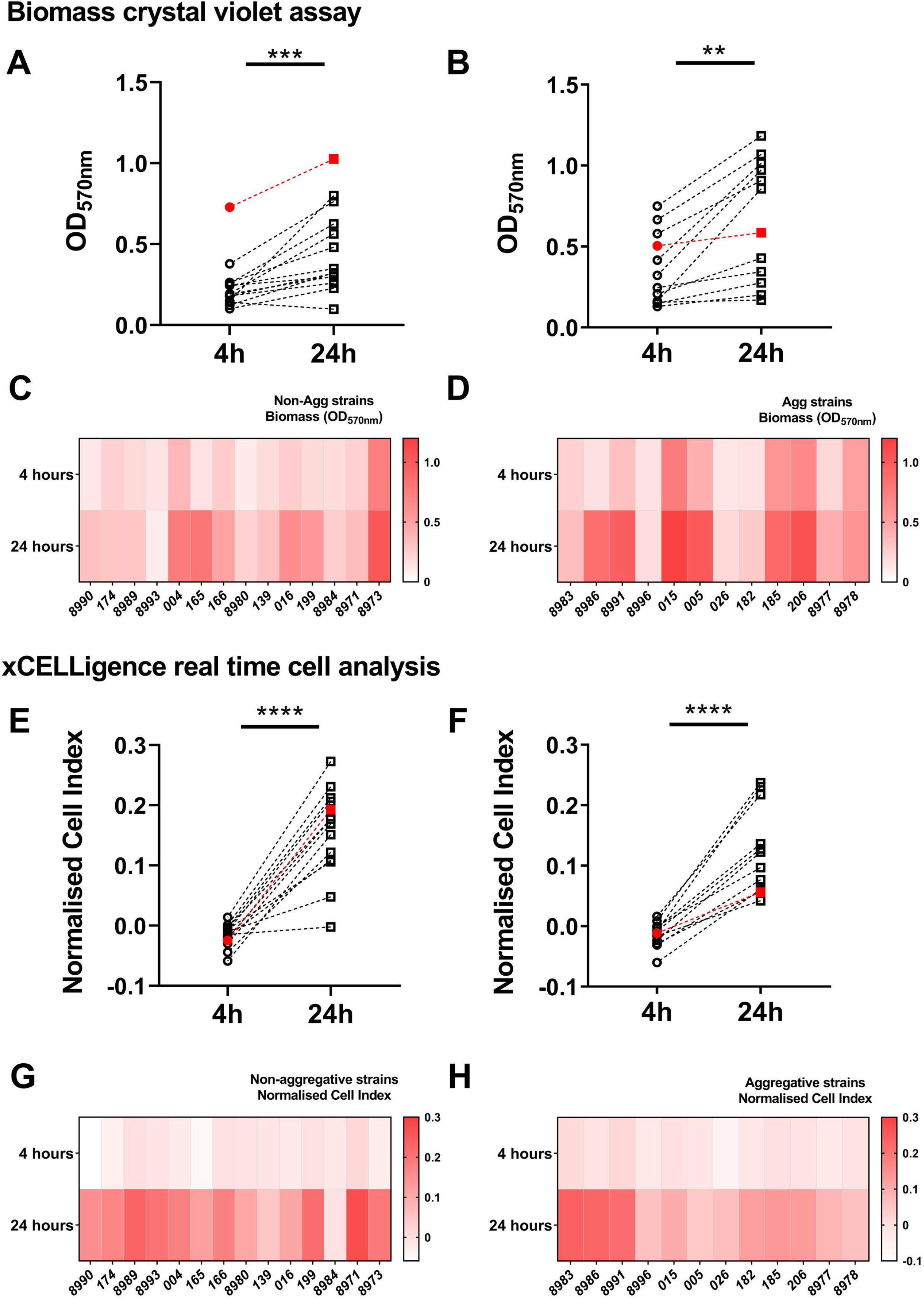
Non-aggregative and aggregative *Candida auris* biofilm heterogeneity. Biomass and impedance measurements were used as measures of biofilm formation of 26 isolates of C. auris (n=14 for non-aggregative and n=12 for aggregative phenotype). For biomass assessment, 1 × 10^6^ fungal cells were seeded in 96-well plates and biofilm developed for 4 h or 24 h prior to crystal violet staining. Panels A and B show the differences in absorbance at 570nm of the non-aggregative and aggregative, at 4 h and 24 h, respectively. The heatmap shows the average absorbance at 570nm for each individual isolate at both timepoints (C and D). The formation of *C. auris* biofilms in real time was monitored using electron impedance measurements on the xCELLigence real time cell analyser. Electron impedance measurements are presented as cell index for all isolates, for biofilms developed for 4 h and 24 h, respectively (E and F). The heatmap depicts the mean cell index values for all non-aggregative (G) and aggregative (H) isolates. Red data points indicate the two isolates selected for further analyses in this study (NCPF 8973 and NCPF 8978). Paired Student’s t-tests were used for statistical analyses and significance between data determined at * p < 0.05 (** p < 0.01, *** p < 0.001, **** p < 0.0001).

To further study the pathogenic and biofilm-forming characteristics of Agg and non-Agg *C. auris*, transcriptional profiling of 24-hour planktonic vs biofilm phenotypes was performed. For these studies, two clinical isolates from the initial pool tested were selected for analysis (non-Agg NCPF 8973 and Agg NCPF 8978, indicated by the red points in Figure 2). Firstly, we found that a total of 701 genes were upregulated in planktonic and/or biofilm form of Agg compared to the non-Agg phenotype, of which 450 genes were upregulated in the biofilm state (Figure 3A). Conversely, less genes (430) were upregulated in non-Agg *C. auris* in the planktonic, biofilm or both states compared to Agg *C. auris* counterparts, with 190 genes up-regulated in the biofilm form (Figure 3B). In order to understand the functional processes related to differentially expressed genes, a cut-off of 2-fold up-regulation was used for gene ontology (GO) analysis (adjusted P value of < 0.05). Up-regulated genes in non-Agg vs. Agg *C. auris* biofilms normalized to planktonic cell expression involved three functional classes; biological processes (BP), cellular components (CC) and metabolic functions (MF) (Supplementary Figure 2A and B). Interestingly, most genes upregulated in the Agg biofilm belonged to the CC functional class; over 40 genes related to membrane and cell wall constituents were upregulated in the Agg form compared to the non-Agg phenotype (Figure 3C). Several genes associated with fungal cell wall proteins were upregulated in the Agg biofilms including *TSA1*, *ECM33*, *MP65* and *PHR1* (Supplementary Figure 2C). Moreover, included in these 40 genes were members of the *ALS* family of adhesins such as *ALS1*. On the contrary, in the non-Agg biofilms, the greatest changes in expression were observed for genes belonging to functional classes of BP and MF (Supplementary Figure 2C). Only a small number of genes belonging to cellular components were upregulated in the non-Agg compared to Agg biofilm (Figure 3D).

**Figure 3.**
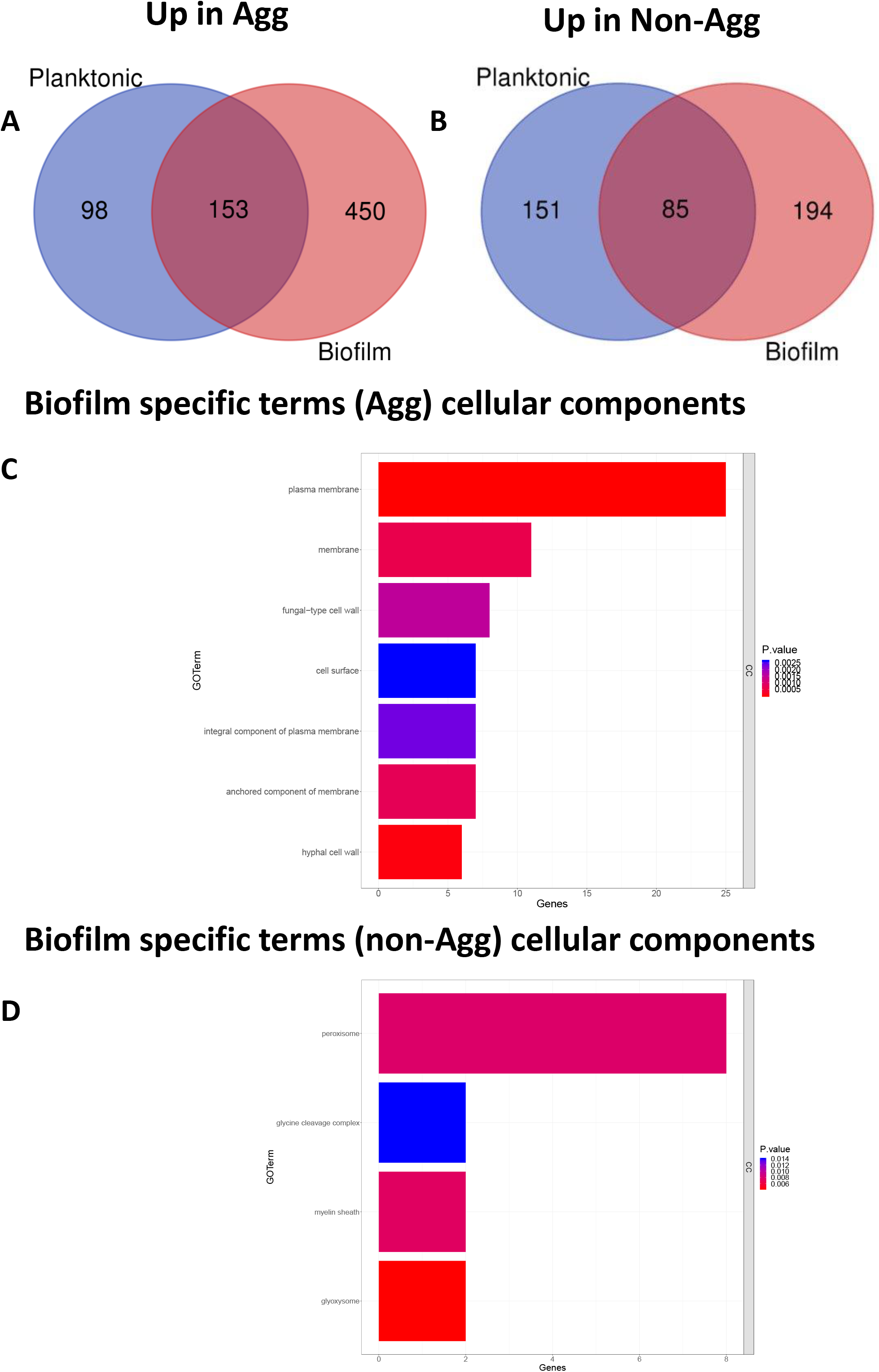
Transcriptional profile of non-aggregative and aggregative *Candida auris* during biofilm formation. RNA from planktonic cells and biofilms formed for 24 h of two isolates (NCPF 8973 and NCPF 8978) was used for RNA sequencing and transcriptome analyses as described in the text. Venn diagrams in panels A and B depict up-regulated genes in non-aggregative (A) or aggregative (B) phenotype in either planktonic form (blue circle), biofilm form (red circle) or in both forms (blue and red circle). In panels C and D, gene distribution of significantly upregulated genes in biofilm forms were grouped for gene ontology analysis. Genes upregulated that belong to the functional pathway cellular components are shown, whilst all other pathways are included in supplementary Figure 2. A cut-off of 2-fold up-regulation was used for gene ontology analysis using an adjusted P value of < 0.05.

The observed differences in the transcriptional profiles of genes belonging to the CC of *C. auris* could impact the ability of the host to recognize the two phenotypes. Key cellular components such as *ALS* proteins of other *Candida* species, such as *Candida albicans* play important roles in aiding host colonization and orchestrating the innate immune response [38, 39]. However, it is currently unknown whether similar mechanisms exist for *C. auris*. Therefore, the following section investigates whether a non-Agg or Agg phenotype dictates the response by the host to *C. auris*. For this, a two- and three-dimensional skin epithelial model was employed to study the host response to the two *C. auris* used above. For both co-culture skin systems, a wound was induced to mimic the possible entry site of patients for *C. auris* in healthcare environments. In the two-dimensional model containing adult human epidermal keratinocytes (HEKa) cells, both isolates were significantly more cytotoxic to host cells following induction of the wound (Figure 4A; **** p < 0.001). Intriguingly, Agg *C. auris* was significantly more cytotoxic than non-Agg form in the wound model (Figure 4A; §§ p < 0.01). It is noteworthy that wounded monolayers minus inoculum were comparable to untreated monolayers (data not shown), suggestive that induction of the wound did not induce cytotoxic effects on the cells. A similar trend in cytotoxicity was observed between the two isolates in the three-dimensional model (Episkin, SkinEthic™ reconstructed human epidermis; RHE) although this did not reach statistical significance (Figure 4B). Interestingly, cytotoxicity of Agg and non-Agg *C. auris* were comparable in both co-culture models minus wounds.

**Figure 4.**
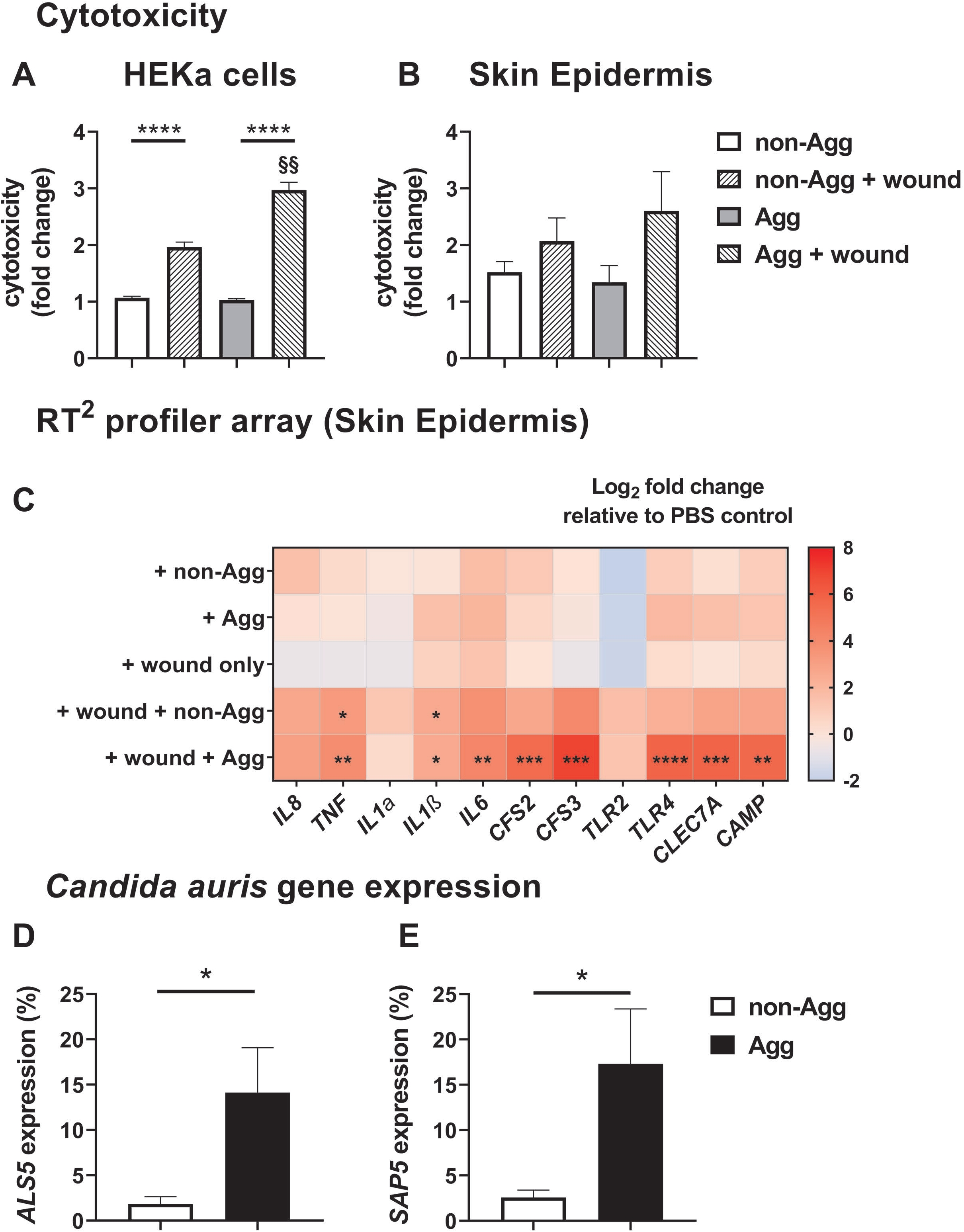
Cytotoxic and inflammatory effects of aggregative and non-aggregative *Candida auris* on skin epithelial models *in vitro*. For these analyses, a two- and three-dimensional skin epithelial model was used as schematically described in Figure 1. Firstly, cytotoxicity in the models were determined by quantifying the amount of lactate dehydrogenase (LDH) released by the human adult epidermal keratinocytes (HEKa) (A) and skin epidermis (B) following co-culture with the aggregative (NCPF 8978) and non-aggregative (NCPF 8973) isolates of *C. auris*. For this, data was presented as fold change relative to PBS control. To study the host response to *C. auris*, a RT^2^ profiler array containing genes associated with inflammation and fungal recognition was utilised to assess the transcriptional profile of the skin epidermis following stimulation (C). Data in the heatmap presented as Log_2_ fold change relative to PBS control. Finally, expression of two virulence genes, *ALS5* and *SAP5* was determined in the isolates infected on the tissue, values presented as % expression relative to the fungal specific house-keeping gene, *β-actin* (D and E). All epithelial cells or tissues were infected in triplicate and statistical significance determined from raw data CT values using unpaired Student’s t-tests for comparison of two variables or one-way ANOVA with Tukey’s multiple comparison post-test for more than two variables (* p < 0.05, ** and §§ p < 0.01, *** p < 0.001, **** p < 0.0001).

To further study the host response in the three-dimensional system, a transcriptional response was investigated using a RT^2^ profiler array containing primers specific for genes associated with inflammatory responses and/or fungal recognition (Figure 4C). Upon investigation, it was evident that the greatest changes in gene expression were observed in the wound models for both *C. auris* isolates. Importantly, induction of the wound minus *C. auris* did not significantly alter the expression of any of the genes arrayed. In the wound model, pro-inflammatory cytokines *TNFα* and *IL1β* (* p<0.05) were significantly upregulated in RHE tissue cultured with non-Agg *C. auris*. Conversely, the Agg phenotype of *C. auris* induced the greatest changes in RHE; the expression of 8 of the 11 genes profiled (*IL1β*, *p<0.05; *TNF*, *IL6*, *CAMP*, ** p<0.01; *CFS2*, *CFS3*, *CLEC7A*, *** p<0.001; *TLR4*, **** p<0.0001) were all significantly upregulated following co-culture compared to the untreated tissue.

In the three-dimensional co-culture system, it was evident from histological and fungal specific Periodic acid-Schiff (PAS) staining that the both isolates of *C. auris* had adhered to the peripheral keratinised layer of the RHE tissue (Supplementary Figure 3). Uninfected tissue displayed a well-organized multi-layered structure characteristic of skin epidermis *in vivo* (Supplementary Figure 3A). However, in infected tissue, there was no sign of *C. auris* infiltration into the tissue (Supplementary Figure 3A and B) as seen with previous publications of skin tissue infection models with invasive *C. albicans* [24, 40]. Unfortunately, loss of tissue integrity in the wounded model rendered them unsuitable for histological staining therefore at this juncture we were unable to visualize whether *C. auris* invaded the tissue at the wound site. Nonetheless, given that the expression of several important cell membrane and wall proteins are differentially regulated in non-Agg and Agg phenotypes of *C. auris* biofilms, we finally wanted to assess the expression of two key virulence factor genes belonging to the ALS and SAP families, respectively. *ALS5* and *SAP5* were both up-regulated in the Agg phenotype compared to the non-Agg *C. auris* isolate in the three-dimensional tissue model (Figure 4D and 4E). Such a response in the Agg isolate may begin to elucidate the mechanism by which this phenotype generated a greater inflammatory response within the tissue.

## Discussion

Results from this study further indicates that the Agg phenotype of *C. auris* determines its pathogenicity *in vitro*. This phenotype, first reported by Borman et al (2016), characterised in certain isolates by the formation of individual yeast cells mixed with large aggregations in planktonic form [6]. This aggregating behaviour was later shown to affect biofilm formation, antifungal susceptibility and virulence of the organism [6, 10, 11, 37]. Moreover, aggregation has recently been found to be an inducible trait triggered by sub-inhibitory concentrations of triazole and echinocandin antifungals, suggestive that treatment regimens must be carefully considered to combat *C. auris* dependant on its Agg phenotype [8]. In this study, we report a previously described RNA-sequencing approach [30] in order to compare the transcriptional responses of one non-Agg (NCPF 8973) and one Agg (NPCF 8978) isolate of *C. auris* during formation of biofilms from planktonic cells. These analyses indicated that several key cell membrane and cell wall components were upregulated in Agg biofilms, many of which involved in cell adhesion to abiotic surfaces and host cells. These observations led us to pose the question: how would the host respond to the two phenotypes? As such, we document for the first time an investigation into the host skin epithelial response to Agg and non-Agg *C. auris*.

Given the clear differences in virulence traits of the non-Agg and Agg phenotype, we deemed it pertinent to study in greater detail the transcriptional profiles of one non-Agg and one Agg *C. auris*. These two isolates displayed the heterogeneity exhibited by other Agg and non-Agg isolates, with the NCPF 8973 isolate forming a denser biofilm after 24 h (as shown here in Figure 2 by the red data points, and elsewhere [10]). However, irrespective of the biofilm-forming capabilities, transcriptome analyses showed that an increased number of genes were upregulated in the Agg compared to the non-Agg biofilms (450 vs 194 genes, respectively; Figure 3), suggestive that the clustering of aggregated cells greatly impacts the transcriptome of the organism during the formation of a biofilm. Of these differential responses in the two biofilm phenotypes, a vast number of genes upregulated in the Agg biofilm belonged to CC as assessed using GO analyses Specifically, several cell wall genes were upregulated in the Agg phenotype e.g., *TSA1*, *ECM33*, *MP65*, *ALS1* and *PHR1*. In *C. albicans*, ECM33 and MP65 have been implemented as important proteins in maintenance of fungal cell wall integrity, biofilm formation and stress responses [41–43]. Furthermore, the ALS family of adhesin proteins and PHR family of extracellular transglycosylases also function as important regulators of biofilm formation in *C. albicans* [44, 45]. For example, loss of function of PHR and ALS proteins results in impaired adhesion and biofilm development [46, 47]. It must be noted here that such comparisons between findings on *C. auris* and *C. albicans* biofilms must be taken with a certain degree of caution given the lack of true hyphal formation of *C. auris* isolates grown in biofilms [6], although such a trait can be induced under certain conditions [7, 14]. Nevertheless, such in-depth analyses, as those documented herein, may begin to explain the mechanisms behind aggregate formation in biofilms of some *C. auris* isolates.

Most of the aforementioned genes (*TSA1*, *ECM33*, *MP65*, *ALS1* and *PHR1*) have multiple functions in *C. albicans* pathogenicity, particularly in biofilm formation as discussed above. In addition, most also play key roles in attachment and/or survival within the host in other fungal species. For example, *TSA1*, which encodes for a protein called thiol-specific antioxidant 1, has been identified in *C. albicans* and *Cryptococcus neoformans*, and functions as an important stress response regulator in unfavourable oxidative environments [48, 49]; potentially those generated by the host[50]. In *C. albicans*, *ECM33*, *MP65* and *PHR1* have been shown to be important genes necessary for production of proteins involved in adherence and invasion of host cells [41–43, 46]. For example, in similar three-dimensional reconstituted skin and oral epithelial co-culture models, heterozygous mutants of *ECM33* and *PHR1* displayed clear deficiencies in penetration of epithelial cell layers leading to reduced tissue invasion and subsequent cellular damage [42, 46]. Finally, another gene upregulated in Agg *C. auris* biofilms was *ALS1*, a member of the ALS family of adhesin proteins. This cohort of adhesin proteins which contains at least 8 members in *C. albicans* are well documented virulence factors, particularly in host-pathogen interactions *in vitro* and *in vivo* [44, 51, 52]. Concerning the results shown in this study, the role of *ALS1,* which encodes for the protein Als1p, in *Candida*-host interactions remains unclear. As such, contradictory reports state a role for this adhesion in attachment to epithelial cells. Kamai and colleagues (2002) showed that heterozygous knockouts of *ALS1* were unable to colonize oral tissues of mice *in vivo* and *ex vivo* as efficiently as wildtype strains or knockouts coupled with the Als1p reinstated [53]. On the contrary, it was shown that attachment of the *C. albicans ALS1* null mutant to oral epithelial cells was no different than wildtype controls, suggestive its role in adhesion was not as important as other *ALS* family members [54]. Nevertheless, as discussed briefly above, such comparisons between *C. auris* and *C. albicans* interactions with host cells must be tentatively compared given the differences in the morphological forms of the two species. In particular, the transition from yeast to hyphae in *C. albicans* is essential for tissue invasion of mucosal surfaces. For example, fungal invasion mechanisms have been identified that show morphological changes from yeast, to pseudohyphae, to hyphae during the process of adhesion and invasion in *C. albicans* in epithelial tissue [55, 56]. This process is even essential for the discrimination of commensal and pathogenic forms of *C. albicans* by the host [57]. Given its inability to form hyphae under normal physiological conditions, how *C. auris* can invade epithelial tissue is unknown. It is possible that the organism has adapted slightly different invasive mechanisms. However, at this juncture, further studies are necessary to elucidate such mechanisms.

It has been postulated that *C. auris* exhibits a level of immune evasion to bypass our immunological defences [4]. Recent work by Johnson et al (2019) showed that *C. auris* were resistant to neutrophil-mediated killing by failing to stimulate neutrophil elastase trap (NET) formation both *in vitro* and *in vivo* in a zebrafish infection model [16]. Another study found that viable *C. auris* did not induce an inflammatory response in human peripheral blood mononuclear cells (PBMCs), although fungal cells were recognized and engulfed by human monocyte-derived macrophages.

Conversely, such host responses were significantly greater against other *Candida* species such as *Candida tropicalis, Candida guilliermondii* and *Candida krusei* [58]. It is apparent from these studies that the host recognizes yet fails to generate an effective immune response against *C. auris*. From the results described here, it is evident that *C. auris* is not cytotoxic nor pro-inflammatory to intact skin epithelial cells or epidermis tissue. Only following induction of a wound in these models did *C. auris* elicit any significant response by the host. From this, it could be suggested that under normal immunological conditions, the organism is non-invasive, and any immune response from the host is minimal. These postulations have been confirmed elsewhere, whereby immunocompetent mice were more resistant to *C. auris* infection than immunocompromised mice [15]. Such findings have been seen in humans; invasive *C. auris* infections generally occur in critically ill patients with serious underlying medical conditions resulting in haematological deficiencies and/or immunosuppression [59–61]. In the context of this study, the observed non-invasive phenotype of *C. auris* may simply be due to lack of hyphal formation, whereby yeast cells colonize the periphery of the skin yet do not invade the underlying layers unless in the presence of a wound. Indeed, similar observations have been made elsewhere. Horton et al (2020) recently showed that *C. auris* forms layers of cells on the periphery of porcine skin *ex vivo*, however, these structures of *C. auris* are devoid of pseudohyphae or hyphae [17].

A limitation from previous studies is the failure investigate the host response to the Agg phenotype expressed by certain isolates. We and others have previously shown that the non-Agg isolate of *C. auris* are more virulent in *G. mellonella* larvae models than the Agg counterparts, possibly resulting from enhanced dissemination rates of the single cells [6, 10]. Here, the Agg NCPF 8978 isolate was considerably more cytotoxic and pro-inflammatory than non-Agg NCPF 8973 in the two- and three-dimensional skin models. At this juncture it is unknown why such a response occurs in the host. Murine model of candidiasis have shown that *C. auris* can accumulate in the kidney of the mice in the form of aggregates [7, 12, 13], suggestive that aggregation may occur *in vivo* to enable persistence and survival. However, to date, no studies have investigated the host response to such *C. auris* aggregates *in vivo*. It could simply be that the Agg phenotype generates a cluster of cells with increased pathogenic traits to induce a greater host response than single cells. This was confirmed by the upregulation of two key adhesin and proteinase genes, *ALS5* and *SAP5*, in the Agg NCPF 8978 isolate compared to non-Agg NCPF 8973 in the skin epidermis model. Interestingly, similar observations have been described in biofilm-dispersed single cells vs. aggregates in model bacteria such as *Pseudomonas aeruginosa*. As such, dispersed aggregates of *P. aeruginosa* possess enhanced antibiotic resistance traits, likely due to encapsulation by extracellular matrix, and greater immune evasion techniques over dispersed single cells[62–64]. Conversely, it could be argued that the single cell phenotype of *C. auris* exhibits a level of immune evasion (as postulated elsewhere [16, 65]), which may explain the lack of response by the host to this phenotype. Future studies must continue to investigate the unique Agg phenotype of *C. auris* to fully clarify the organisms’ pathogenic mechanisms. These investigations must consider interactions between *C. auris* and other organisms that comprise the skin and/or wound microbiomes, which may function as important beacons for host invasion of *C. auris*.

The identification of five geographically and phylogenetically distinct clades of *C. auris* [2, 9], containing isolates capable of forming aggregates with enhanced drug resistance, has meant that unravelling the pathogenic mechanisms employed by *C. auris in vitro* and *in vivo* remains extremely difficult. This study has highlighted different pathogenic signatures of Agg and non-Agg forms of *C. auris* in biofilms and during host invasion *in vitro*. Of course, given the level of heterogeneity amongst isolates, in-depth analyses from this work are limited to one Agg and one-Agg isolate. As such, future studies must continue to investigate these unique phenotypic traits of different Agg and non-Agg isolates of *C. auris* to fully understand the persistence of this nosocomial pathogen in the healthcare environment, and whether such traits are comparable amongst the diverse isolates.

## Acknowledgements

The authors would like to thank the Glasgow Imaging Facility (University of Glasgow) and Margaret Mullin for assistance in scanning electron microscopic techniques. The authors would like to acknowledge funding support of the BBSRC Industrial GlaxoSmithKline CASE PhD studentship for Christopher Delaney (BB/P504567/1).

## Declaration of interest

The authors declare no conflicts of interest.

